# multiPhATE: bioinformatics pipeline for functional annotation of phage isolates

**DOI:** 10.1101/551010

**Authors:** Carol L. Ecale Zhou, Stephanie Malfatti, Jeffrey Kimbrel, Casandra Philipson, Katelyn McNair, Theron Hamilton, Robert Edwards, Brian Souza

## Abstract

**Summary:** To address the need for improved phage annotation tools that scale, we created an automated throughput annotation pipeline: multiple-genome Phage Annotation Toolkit and Evaluator (multiPhATE). multiPhATE is a throughput pipeline driver that invokes an annotation pipeline (PhATE) across a user-specified set of phage genomes. This tool incorporates a *de novo* phage gene-calling algorithm and assigns putative functions to gene calls using protein-, virus-, and phage-centric databases. multiPhATE’s modular construction allows the user to implement all or any portion of the analyses by acquiring local instances of the desired databases and specifying the desired analyses in a configuration file. We demonstrate multiPhATE by annotating two newly sequenced *Yersinia pestis* phage genomes. Within multiPhATE, the PhATE processing pipeline can be readily implemented across multiple processors, making it adaptable for throughput sequencing projects. Software documentation assists the user in configuring the system.

**Availability and implementation:** multiPhATE was implemented in Python 3.7, and runs as a command-line code under Linux or Unix. multiPhATE is freely available under an open-source BSD3 license from https://github.com/carolzhou/multiPhATE. Instructions for acquiring the databases and third-party codes used by multiPhATE are included in the distribution README file. Users may report bugs by submitting to the github issues page associated with the multiPhATE distribution.

**Supplementary information:** Data generated during the current study are included as supplementary files available for download at https://github.com/carolzhou/PhATE_docs.

## 1 INTRODUCTION

Global pathogen discovery efforts, such as The Global Virome Project (Carrol et al. 2018), are projected to invest billions of dollars to support surveillance projects that characterize the earth’s virosphere over the next 10 years. Already, the PhagesDB contains more than 13,000 phage genomes (Russell et al., 2017). Phage therapy has resurfaced as a method to combat antimicrobial resistance, and upcoming clinical trials necessitate complete sequencing and characterization of therapeutic candidates, but high-quality gene finding and functional annotation are vital for successful genomic comparison studies and for discovery of new phage-based therapeutic leads (Kutter et al., 2015). Because annotation of phage genomes is a relatively new science, there exist few bioinformatics pipelines for phage analysis that can be readily adapted for use in phage research efforts. Currently, researchers typically apply bacterial gene finders for annotation of phage DNA, followed by largely manual analyses using web forms, and integration of summary results can be time consuming. Although there exist several codes for identifying prophage sequences in bacterial genomes (Arndt et al., 2016; Kang et al., 2018; Roux et al., 2015; and others), once these sequences have been identified, they are typically annotated using methods developed for sequences from other taxa (Perkel, 2017; Seemann, 2014). Currently there exists only one automated annotation pipeline specifically for phage: Philipson et al., 2018, describe a pipeline that identifies features in phage that determine their potential suitability as therapeutic reagents. However, there remains a need for an automated phage annotation pipeline that can be readily implemented on multiple nodes of a local server and that requires minimal software development expertise. To address this need, we present the multiPhATE automated phage annotation pipeline.

## 2 DESCRIPTION

The PhATE annotation pipeline incorporates four gene finders: GeneMarkS (Lomasadze et al., 2017), Glimmer (Delcher et al., 2007), Prodigal (Hyatt et al., 2010), and a novel phage-centric gene finder, THEA (McNair et al., doi: https://doi.org/10.1101/265983) (now called PHANOTATE). Functional annotation is achieved by BLAST and HMM searches for homologous sequences in protein-and phage-centric databases. The PhATE workflow is depicted in suppl. file, “phate_Fig_1_PhATE_Workflow.pdf”. **Input.** Input to multiPhATE consists of a configuration file that specifies a list of genomes to be processed by PhATE and a set of parameters controlling software execution. The user specifies the names of phage genome fasta files, the names of output subdirectories, and other metadata pertaining to the genomes being analyzed. The user also specifies the following optional analyses: (a) gene caller(s) to be run; (b) gene-caller to use for subsequent annotation (default: PHANOTATE); (c) blast parameters; (d) blast databases to be searched; (e) turn hmm search on/off. It is possible to run PhATE using any or all of the specified gene callers, databases and searches. In this way, installation can be achieved one gene-caller or database at a time, with stepwise testing. Also, the user can switch on/off searches (e.g., NR) in order to control execution time (this may be useful in performing preliminary annotation of large numbers of sequences). Although multiPhATE is intended for phage sequence annotation, it would be reasonable to run multiPhATE with bacterial genomes to assist identification of embedded phage sequence.

### Annotation

PhATE begins by performing gene calling using the selected gene finder(s). When two or more are invoked, PhATE outputs a summary table showing a side-by-side comparison of the gene calls, plus summary statistics regarding the numbers and lengths of gene calls for each algorithm, and the numbers of calls in common and unique to each. Next, PhATE uses BLAST+ programs (Camacho et al., 2008) blastn and blastp, and the HMM search program jackhmmer (Johnson et al., 2010), to identify homologs of the input genome and its predicted gene and peptide sequences using several databases: NCBI virus genomes, NCBI Refseq proteins, NCBI refseq genes, NCBI virus proteins, and NR (NCBI Resource Coordinators, 2016), as well as Swissprot (Bairoch and Appweiler, 2000), PhAnToMe (www.phantome.org), a virus subset of KEGG (Kanehisa et al., 2017), and a fasta sequence data set derived using the pVOG identifiers (Grazziotin et al., 2017). The latter database is modified to contain the pVOG identifiers in the fasta headers, by means of scripts included in the multiPhATE distribution.

### Output

PhATE generates the following files and directories: (a) output from the gene-call algorithms and the gene-call comparison (suppl. file: phate_P2_CGC.pdf); (b) gene and translated peptide fasta files; (c) combined-annotation summary files, (d) directories containing raw BLAST outputs for genome and peptide blast runs; (e) directories with raw HMM search outputs for peptide searches; (f) alignment-ready fasta files containing each predicted peptide plus the members of each identified pVOG family to which a peptide may be assigned; (g) log files. BLAST and HMM raw data outputs can be saved or cleaned from the output directories (see README). We demonstrate application of multiPhATE to the annotation of two newly sequenced *Y. pestis* phage genomes (see suppl. file, “phate_results.pdf”.

## ACKNOWLEDGEMENTS

This work was performed under the auspices of the US Dept. of Energy by Lawrence Livermore National Laboratory under contract DE-AC52-07NA27344. See suppl. file, “phate_author_information.pdf”.

